# Chromatin landscape and enhancer-gene interaction differences between three cardiac cell types

**DOI:** 10.1101/2025.09.23.678180

**Authors:** Yonglin Zhu, Jean-Christophe Grenier, Raphaël Poujol, Svenja Koslowski, Olivier Tastet, Chang Jie Mick Lee, Matthew Ackers-Johnson, Roger Foo, Julie Hussin, Chukwuemeka George Onyeka Anene-Nzelu

## Abstract

Genome-wide association studies (GWAS) have identified numerous single nucleotide polymorphisms (SNP) associated with a specific traits and diseases, however, uncovering the true disease-relevant SNPs remains challenging. One limitation for prioritizing true disease-relevant SNPs from GWAS is that most of the identified SNPs are non-coding, making it difficult to unravel their mechanism of action. Nevertheless, mapping non-coding SNPs to enhancers is a validated approach to link SNPs to their target genes through the analysis of enhancer-gene interactions (EGI) and thus provide insight into their mechanism of action. While previous studies linking cardiac disease-relevant SNPs to enhancers and their target genes have focused on the principal cardiac cell type, cardiomyocytes (CMs), the analysis of other non-CM cell types has been largely ignored and has only gained attention recently. We hypothesize that characterizing cell-type-specific enhancer-gene interactions (EGIs) for these non-CMs, namely cardiac fibroblasts (CFs), endothelial cells (ECs), and smooth muscle cells (SMCs), followed by mapping cardiac-disease-associated non-coding SNPs to those enhancers will identify novel disease-relevant genes and provide insights for future mechanistic research. To identify the landscape of cell-type-specific EGIs in these cardiac cells, we have employed the activity-by-Contact (ABC) model. It integrates assay for transposase-accessible chromatin sequencing (ATAC-seq), H3K27ac chromatin immunoprecipitation with sequencing (ChIP-seq), and high-throughput chromosome conformation capture with H3K27ac immunoprecipitation (H3K27ac HiChIP) data to identify EGIs. We have identified the landscape of cell-type-specific EGIs in these cardiac cells. Furthermore, a higher similarity of the chromatin accessibility profile (ATAC-seq) between CF and SMC, compared to CF and EC, and SMC and EC was observed. Finally, overlapping identified EGIs with cardiac-disease-associated non-coding variants has allowed the identification of a QT-interval-associated SNP that is mapped to the enhancer region of an EC-specific EGI.

## INTRODUCTION

Genome-wide association studies (GWAS) aim to associate variations in the human genome sequences to specific diseases or phenotypes in the hope of identifying disease/trait-relevant genes ^1^. These studies have increased our understanding of the genomic architecture of diseases, facilitated the identification of genomic risk factors in patients, and assisted in the development of novel therapies. In a typical ‘cases vs. controls’ GWAS analysis, hundreds to thousands of genomic variants are compared simultaneously to identify those variants associated with the identified trait in a statistically significant manner (commonly p < 5 *×* 10^−8^) ^2,3^. The most common genomic variants identified through GWAS are single-nucleotide polymorphisms (SNPs) ^1^. However, identifying those variants itself does not necessarily provide any information on disease-associated genes or mechanisms. Indeed, more than 80 percent of the variants are identified in non-coding regions of the genome, such as putative regulatory elements ^3,4^. Further, a reported disease-associated SNP might not be the causal variant ^5,6^. This can happen due to linkage disequilibrium (LD), where nearby SNPs that are inherited together, only one of them has been reported in the original GWAS. It is therefore essential to consider the SNPs in LD with the GWAS-reported variant. Disease-relevant gene identification and mechanistical validation of these non-coding variants is still only in its infancy, and it will demand substantial scientific and economic effort before therapeutic applications can reach patients.

A recent review has pointed out the importance of enhancer and enhancer variant characterization for the interpretation of GWAS data ^7^. Enhancers are crucial DNA sequences that are typically 50 bp to 1.5k bp in size. They regulate gene transcription regardless of their orientation and distance from their target gene or genes ^8^. Functionally, enhancers stabilize the transcription machinery, thus sustaining the expression of their target genes, and increasing the amount of the corresponding gene product ^9,10^. Thus, they play a crucial role in gene expression regulation to carry out context-dependent roles for proper embryonic development, cell type differentiation, and environmental feedback ^9–11^. Interestingly, the estimated number of enhancers in the human genome is higher than the number of coding genes ^12^, and indeed, a single genes expression has been known to be modulated by more than one enhancer to account for different stimuli, developmental stages, etc ^11^. On the other hand, enhancers are not gene-specific like promoters since they may regulate the expression of more than one target gene ^8^. The context-specificity of enhancer-gene interactions (EGIs) can therefore confound the identification of relevant target genes. Characterizing cell-type-specific enhancer-gene interactions (EGIs) can thus help increase not only our understanding of cell identity, but also decipher the contributions different cell types may play in disease development. Finally, this knowledge will assist the development of more targeted treatment strategies ^13^.

Chromatin accessibility, which describes DNA-binding availability by transcription factors (TFs), can be characterized into the more accessible euchromatin and the less accessible, hence inactive heterochromatin. It is subject to change by signals like biological prompts, environmental cues, and disease states ^14^. Methods like the assay for transposase-accessible chromatin with sequencing (ATAC-Seq) ^15,16^ allow to characterize chromatin accessibility genome-wide, allowing researchers to gain insight into which genomic regions could be transcriptionally active.

Post-translational modifications of histone tails, such as acetylation and methylation ^17^, can modulate genome condensation and, therefore, heterochromatin and euchromatin states. Histone acetylation, one of the most widely studied histone modifications ^18^, can decrease the electrostatic attraction between the negatively charged DNA phosphate backbone and the positively charged histone tail, therefore increasing the accessibility of DNA to other DNA-binding proteins, such as transcription factors (TF). Hence, histone acetylation is generally considered as a marker of active gene expression ^19^. A well-known example of histone acetylation is H3K27ac (acetylation of lysine 27 of the H3 histone subunit) ^18^, and it is one of the most studied active enhancer markers ^19–21^. Chromatin immunoprecipitation with sequencing (ChIP-seq) combines immunoprecipitation and next-generation sequencing (NSG) techniques to specifically identify DNA regions that interact with proteins (e.g. TFs) or histone modifications of interest ^22–24^. Thus, in combination with chromatin accessibility assessment through ATAC-seq, H3K27ac ChIP-seq allows to determine active enhancer regions. However, those assays do not provide any information about the corresponding target gene(s) of an enhancer. Chromatin conformation capture techniques like Hi-C ^25^ and HiChIP ^26^ allow to characterize the interactions between two DNA segments, for example, an enhancer to its target gene. By crosslinking spatially proximate DNA segments in the nucleus before sequencing the 3D genomic architecture can be depicted. However, physicial interaction alone does not prove the activity of those segments.

Thanks to advances in next-generation sequencing and bioinformatic software, an increasing number of tools have been developed to unravel the EGI. One of such methods is called activity-by-contact (ABC) ^27^. ABC integrates the chromatin landscape sequencing data, which includes chromatin accessibility (ATAC-seq), histone modifications (ChIP-seq), and 3D chromatin architecture (HiChIP) ^18^ and computes a score for each identified EGI. The score, which could be intepreted as the percentage contribution the enhancer has to regulate a given gene’s expression, is related to the strength of activating chromatin marks (enhancer activity) and physical contact freuqency between the enhancer and the gene (contact). The activity of the enhancer is estimated from chromatin accessibility (ATAC-seq) and H3K27ac histone modification sequencing (ChIP-seq) data, while the target gene and contact frequency between the two DNA segments is estimated from 3D chromatin architecture (HiChIP). These tools have deepened our understanding in the different gene expression patterns between different cell types, which can help us elucidate the contribution of different cell types in complex diseases like cardiac disease.

Cardiac disease (CD) is the second leading cause of death in Canada ^28^, and numerous GWAS studies have been carried out in recent decades to decipher underlying genetic risk factors. As cardiomyocytes (CMs) represent the predominant and contractile cardiac cell type they have been extensively studied in the context of CD ^29^. Indeed, a recent study has mapped the EGI landscape of CMs to identify the targets of non-coding CD-associated SNPs ^30^. Here, we have extended this approach to investigate the EGIs in heart-resident non-cardiomyocyte (non-CM) cell types, namely cardiac fibroblasts (CF), endothelial cells (EC), and smooth muscle cells (SMC), which are three of the most abundant non-CMs ^29^. These cells play crucial supporting roles for the proper functioning of CMs and the heart as a whole. Each of them expresses different sets of genes to perform their necessary functions ^31^, even though they originate from the same layer during embryonic development ^32,33^.

Even though in recent years EGIs are increasingly studied, a comprehensive map of EGIs in non-CM cell types of the heart has not been established yet. The characterization of the EGI landscape of all cardiac cell types could contribute valuable information to gain mechanistic insights how genomic variants influence cardiac function and disease. Here we present our work, where we have characterized EGIs in three different non-CM cell types using ATAC-seq, H3K27ac ChIP-seq, and H3K27ac HiChIP data and subsequent ABC-score calculations.

## RESULTS

### Cardiac fibroblasts and human coronary artery smooth muscle cells share more similar chromatin landscapes

Diffbind ^34^ identified 152,455 open chromatin sites across 11 samples, with 103,744 to 118,604 peaks in cardiac fibroblast (CF) samples, 63,601 to 79,073 in human coronary artery endothelial cells (HCAEC), and 89,272 to 93,363 in human coronary artery smooth muscle (HCASMC) cells. Roughly 20% of all peaks were intergenic, the rest were either in genic regions or promoter regions. Principal component analysis (PCA) (Supplementary fig. S1) and correlation heatmaps of ATAC-peaks (Fig. 2 B) have been generated as quality control before differential analysis.

**Figure 1:**
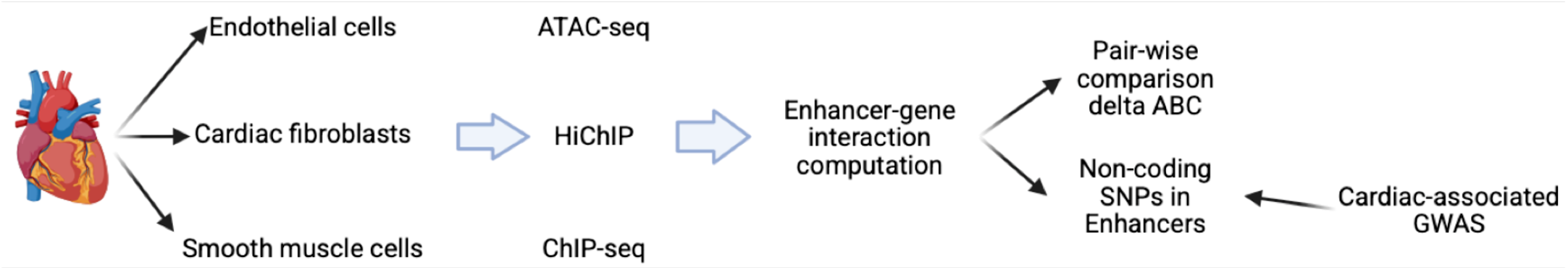
Graphical Abstract.

**Figure 2:**
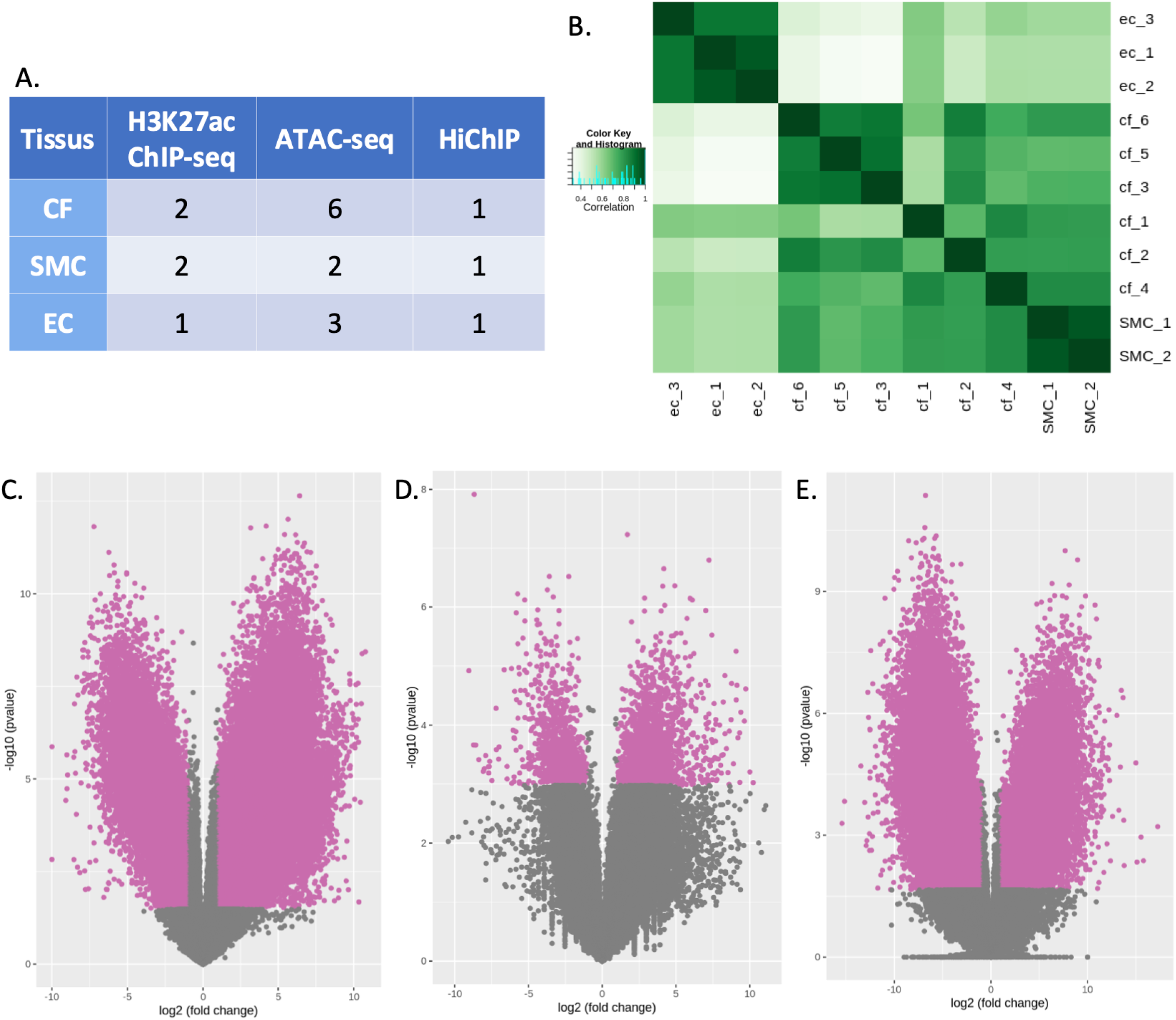
Open chromatin profiles of the three cardiac cell types. A. Summary of number of samples for each sequencing and each cell type. *EC: Endothelial cells; CF: Cardiac fibroblasts; SMC: Smooth muscle cells* B. Sample similarity and correlation of ATAC-seq signals. Sample correlation was performed using the Diffbind package utilizing the Pearson method. The darker the square, the higher the similarities between the two samples. C. - E. Pair-wise comparisons of ATAC-seq of the three cardiac cell types. Each dot represents an open chromatin region or a single ATAC-seq signal. The pink dots represents regions that are have False discovery rate (FDR) < 0.05 and absolute log2 fold change > 1. C. Volcano plot showing endothelial cell (EC) against cardiac fibroblasts (CF). A positive fold change stands for stronger signal in EC. D. Volcano plot showing smooth muscle cell (SMC) against cardiac fibroblasts (CF). A positive fold change stands for stronger signal in SMC. E. Volcano plot showing smooth muscle cell (SMC) against endothelial cell (EC). A positive fold change stands for stronger signal in SMC.

Pairwise comparison of accessible chromatin regions between cell-types were performed (see Methods). Only the regions showing significantly differented accessibility in both, by DESeq2 and edgeR, were retained (Fig. 2 C. - E.). 97,995 (edgeR) to 102,375 (DESeq2) peaks were identified as significantly different regions between CF and HCAEC, using CF peaks as the reference; 65% of all the open chromatin regions of HCAEC and CF were different. For CF and HCASMC, 3,167 (edgeR) to 13,166 (DESeq2) peaks have been identified as significantly different using CF as reference, 2% of all the open chromatin regions differed significantly between those two cell-types. Between HACEC and HCASMC, 66,735 (edgeR) and 73,918 (DESeq2) peaks were significantly different, using HCAEC as references, which was about 44% of all the open chromatin regions.

As expected, differences among the chromatin accessibility profiles of the three non-CM cell-types was observed. The total number of accessible chromatin regions were roughly comparable between the three cell types (Supplementary table. S2).

Interestingly, we observed a higher similarity between HCASMC and CF than between CF and HCAEC and HCASMC and HCAEC, suggesting a higher overlap in active genomic regions between HCASMC and CF than between either of those cell-types (Fig. S1). For H3K27ac ChIP-seq, 82,310 to 84,786 peaks were identified in cardiac fibroblast samples, 43,962 peaks in HCAECs, and 65,338 to 73,294 peaks in HCASMCs. Pairwise comparison of differential H3K27ac ChIP-seq peaks between HCASMC and CF seems to support the observation of similarity in the active genomic regions between the two cell types. Indeed, only 5,295 and 814 significantly different peaks were identified by edgeR and DESeq2 between the two, respectively.

### The Activity-by-contact (ABC) model identifies/highlights different enhancer-gene interactions (EGIs) between non-CMs

To account for individual sample and sex variations, we computed a consensus ATAC peak set for each cell type. MACS2 was able to identify 118,300 (CFs), 64,233 (HCAECs), and 80,184 (HCASMCs) consensus ATAC peaks that have been identified in at least 2 samples for each of the three cell types. These consensus peak sets were used as the ATAC-seq input files for the ABC model, which in combination with H3K27ac histone modification signals, have been used to identify putative enhancers for the three cell types (Table S3).

Through the integration of the ATAC-seq, H3K27ac ChIP-seq and H3K27ac HiChIP data for each cell-type, the ABC model identified 8,366,654, 5,561,019, and 6,248,721 putative EGIs in total in CF, HCAEC and HCASMC, respectively. The total number of putative interactions were comparable between the three cardiac cell types.

### Shared EGIs between two cell types (Common EGIs)

We next searched for the common EGIs between each two of the three cell types. We included all of the EGIs identified by the ABC, even those below the 0.05 cut-off, to culminate the maximum number of common EGIs. Indeed, an EGI that is biologically irrelevant for the regulation of a given gene (ABC score < 0.05) in one cell type can be an important regulator of the gene’s expression in a different cell type (ABC score > 0.05). 3,589,912 common EGIs have been identified between HCAECs and CFs, 4,807,359 between HCASMCs and CFs, and 3,599,150 between HCAECs and HCASMCs. Interestingly, the high number of common EGIs for HCASMCs and CFs reflects our previously observed similiarity between these two cell types and strengthens the hypothesis that similar gene expression patterns may be active.

We next calculated the ABC score differences (delta ABC) for each of these common EGIs. We have chosen a threshold for significant delta ABC of mean *±* 5 *×* standard deviation of the delta ABC distribution for that overlap, based on the overall distribution of all delta ABC scores. We observed that most of the delta ABCs values are close to zero, indicating that a majority of enhancers contribute to a similar level to the expression of a given gene in both cell types (Figure 3), which was somewhat expected as all of these three cell types were of heart origin, therefore sharing lots of similarities in gene expression. However, as displayed in that figure, this method also allowed us to identify a number of highly cell-type specific EGIs for each comparison.

**Figure 3:**
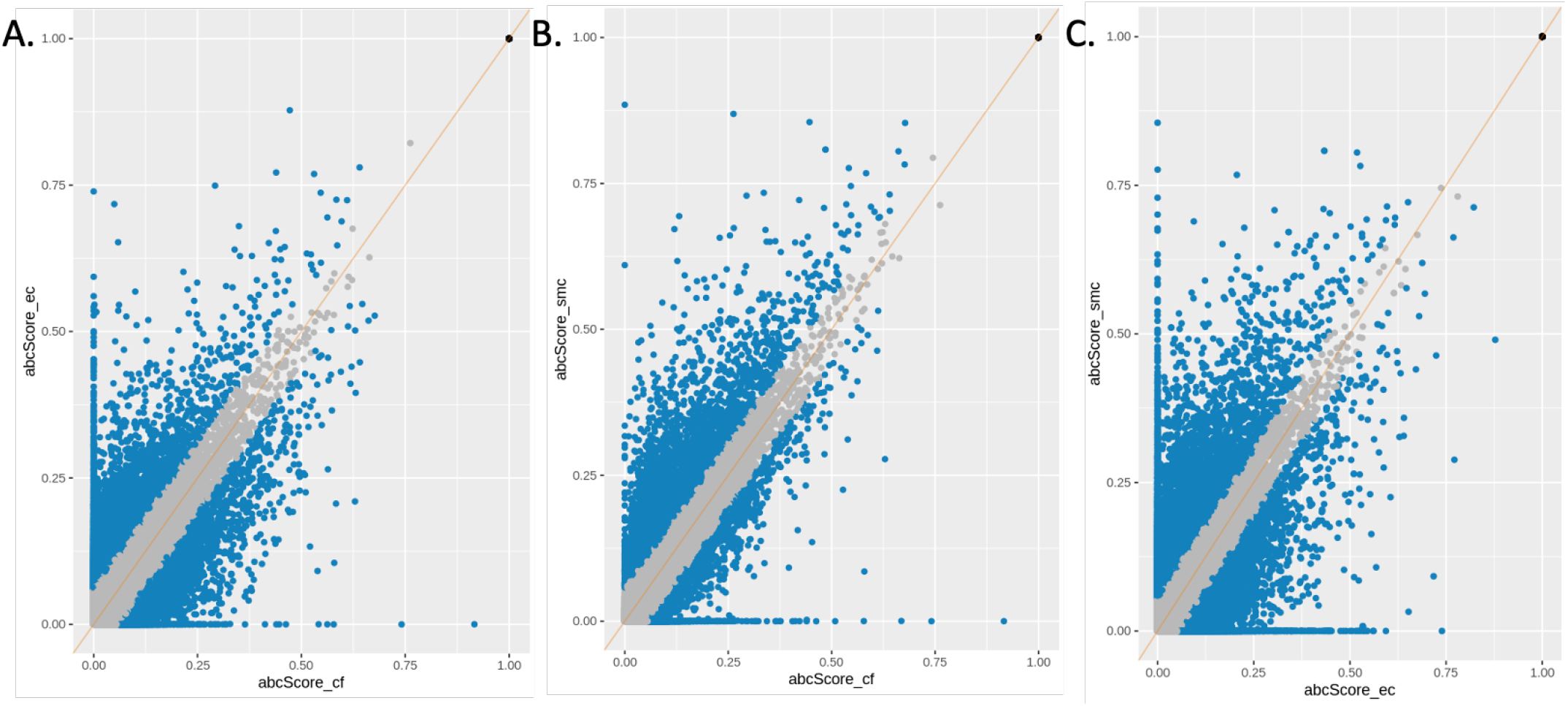
Pair-wise common enhancer-gene interactions (EGIs) and abc score distribution. Each point represents an EGI that have been in both cell types on a scatter plot with each axis representing the ABC score of a cell type. The grey dots represent the EGIs with a delta ABC less than the threshold (mean *±* 5 *×* standard deviation of the delta ABC distribution), while the blue ones are the EGIs that have significant differential delta ABC in the two corresponding cell types. The black dots represents promoters where their ABC scores in both cell types are 1. A. Scatter plot showing endothelial cell (EC) against cardiac fibroblasts (CF). X-axis: ABC score for CF, y-asix: ABC score for EC. B. Scatter plot showing smooth muscle cell (SMC) against cardiac fibroblasts (CF). X-axis: ABC score for CF, y-asix: ABC score for SMC. C. Scatter plot showing smooth muscle cell (SMC) against endothelial cell (EC). X-axis: ABC score for EC, y-asix: ABC score for SMC.

We then filtered out the common EGIs for three comparisons, which left us the cell-type-specific EGIs for these three non-CMs. In total, we have identified 3,046,142, 1,454,505, and 949,901 cell-type-specific EGIs for CF, HCAEC, and HCASMC, respectively.

### Cardiac phenotype-associated SNPs that fall in the enhancer region of cell-type-specific EGIs

Finally, for all the EGIs, we were interested in whether they were involved in cardiac-disease phenotypes. Therefore, we decided to investigate whether single nucleotide polymorphisms associated with a cardiac phenotypes in GWAS studies (GWAS catalogue ^35^) overlapped with the enhancer regions of our EGIs. In total, we mapped more than 68,000 SNPs (sentinel SNPs and SNPs in linkage disequilibrium (LD) with the sentinels) to all identified EGIs with the aim to identify variants situated in cell-type specific enhancers. Among the enhancer regions that host a higher number of non-coding cardiac-phenotype-associated SNPs, we singled out an intergenic region from chromosome 7 (genomic coordinates of hg38 chr7:150988285-150988785). This region contains a SNP (rs5888424) that has not been extensively studied yet. However, this SNP has been found in LD with three other cardiac sentinel SNPs (rsID: rs2968863, rs2968864, and rs1805123), all of which were associated with QT-interval phenotypes ^35^. The enhancer interacts with the NOS3 gene (nitric oxide synthase 3) based on our ABC computation. The NOS3 gene is a known endothelial gene with high expression in cardiac endothelial cells (Fig. S5) ^36^. The primary function of its gene product is to produce nitric oxide (NO), which is shown to have a protective role in cardiovascular disease ^37^, while the detailed mechanism remains unclear ^38^. Interestingly, NOS3 has also been found to be related to QT-related cardiac disease ^39^. As illustrated in Figure 4 A, this enhancer seems to have a high specificity to endothelial cells, while being negligable (ABC score < 0.05) in cardiac fibroblasts. This observation is supported by our ATAC-seq data, which shows that the chromatin in this enhancer region is much more accessible than in CF (Fig. 4 B). Other examples of genes targeted by enhancers of cell-type-specific EGIs, that also harbour cardiac phenotype-associated SNPs, include NOTCH1 in EC (Fig. S6) and HSPG2 in CF (Fig. S7), which play important roles in their respective cell types ^40,41^.

**Figure 4:**
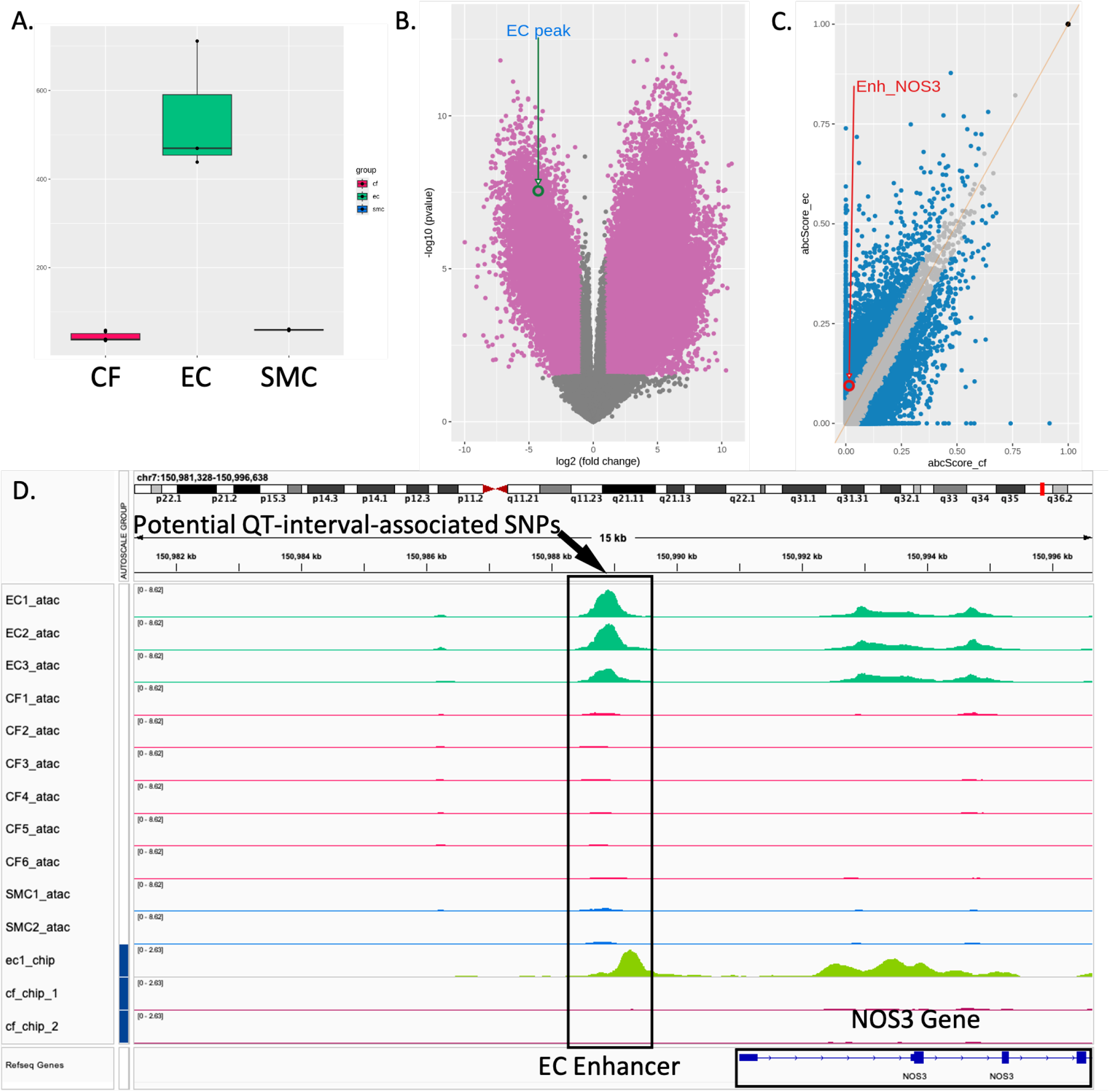
Example of EC-specific enhancer(chr7:150988285-150988785)-NOS3 interaction. Enhancer(chr7:150988285-150988785)-NOS3 interaction that host a QT-interval-associated SNPs in endothelial cells. A. Nomralized counts of chromatin accessibility (ATAC) profile for the enhancer region in all three cardiac cell types. B. The enhancer ATAC signal on the EC vs CF volcano plot. C. The EGI on the pair-wise delta ABC score scatter plot. D. The enhancer(chr7:150988285-150988785)-NOS3 interaction EGI visualized using Integrative Genome Viewer (IGV).

## DISCUSSION

Genome-wide asoociation studies (GWAS) allow us to associate genomic variations, such as SNPs with specific phenotypes like diseases ^1,3^, and hold considerable promise helping us identify disease-relevant genes and pathways. However, the majority of GWAS-identified SNPs lie in the non-coding region of the genome ^42^, which complicates the identification of culprit genes and mechanisms. Resent research has shown that characterising the interactions between a gene and its regulatory element (for example, an enhancer) can help identify the role of non-coding genomic variants ^4,42,43^. As numerous regulatory elements in the genome exercise their function in a cell-type- and/or contex-dependent manner, a better understanding of their specificity is crucial to exploit the full potential of GWAS studies. In this proof-of-concept study, we have focused on the three major non-cardiomyocyte cell populations of the heart, and with our approach identified an HCAEC-specific regulatory SNP in the enhancer region of the NOS3 gene associated with QT-interval phenotypes.

The ABC model we applied defines and quantifies enhancer-gene interactions (EGIs) by integrating H3K27ac histone modification ChIP-seq, ATAC-seq, and chromatin conformation capture data, and it computes a contribution score of an enhancer’s effect on the regulation of a gene’s expression ^27^. Previously, this approach has been shown successful in cardiomyocytes (CM), the major and contractile cell-type of the heart ^30^. Here, we have extended it to three other major cardiac cell types, cardiac fibroblasts (CFs), human coronary artery endothelial cells (HCAECs), and human coronary artery smooth muscle cells (HCASMCs), to investigate the contribution of non-cardiomyocytes (non-CMs) to cardiac disease. For example, CFs, which are important in maintaining heart structure and providing mechanical support by expressing extracellular matrix (ECM) proteins, notably collagens ^29^, are the main effectors for the condition called cardiac fibrosis, where excessive ECM proteins deposit in the heart after cardiac injury or pressure overload ^44,45^. Broadly, SMCs constitute an integral part of the cardiac vessel wall to support blood outflow from the heart ^29^. They can also differentiate to have an expression pattern resembling other cell types in disease states like atherosclerosis ^46^. Moreover, SMCs share functional similarities with CFs, in that they both synthesize and secrete ECM proteins ^29^, even though these two cell types mainly reside in two different regions in cardiac tissues ^47^. Cardiac ECs mainly consist of cells that line the blood vessels, the endocardium, and lymphatic vessels ^29^. Unlike the cardiac SMCs, which make up larger blood vessels, more than 50% of cardiac ECs are capillary ^48^, indicating at their involvement in homeostasis maintenance and immune regulation of the heart. Certain subtypes of ECs have been found to have cardiac function-preserving abilities in coronary artery diseases ^47^.

Overall, vastly different epigenetic profiles and EGIs between CFs, HCAECs, and HCASMCs have been observed. This study established the groundwork that non-CMs, for them to have distinct supporting roles in the heart function, have a different active genomic profile. Understanding the genes in play and their mechanisms, which to this day is still not fully clear, will benefit future cardiovascular therapeutic research.

While individual studies have carried out chromatin accessibility and histone modification analyses in those cell types ^49–51^, this study is the first to compare such profiles between cell-types with the aim to characterize on a genom-wide scale cell-type specificity of EGIs. More specifically, we have identified different open chromatin profiles between these three non-cardiomyocyte cell types. Through our analysis, we have demonstrated a similar profile of chromatin accessibility between CF and HCASMC compared to HCAEC. This suggests that HCASMC and CF have similar profiles of genes activity, or that HCAEC present a more distinct active gene profile than the other two cell types, which calls for future transcriptiomic and proteomic characterizations, such as RNA-seq and western blots. As both cell types share similar functions, such as contribution to extracellular matrix activation and proliferation, and cardiac fibrosis, it raises the question of crosstalk and similarity between CFs and other smooth muscles cell-types ^29^. We do, however, recognize that our data lacks single-cell resolution which would allow to identify potential subtypes of these non-CMs, which could present variable expression and hence open chromatin profile ^45^.

Furthermore, our approach in this proof-of-concept study allows to compare cell-type specific EGIs, as well as the relative contribution of a given enhancer to the expression of a specific target gene. This advantage is conferred by the non-binary output generated by the ABC score method. We have identified a list of cell-type-specifc EGIs and common EGIs shared between at least two of the three non-CM cell types. Our lists of cell-type-specific and significantly differentially active common EGIs is a first step towards future functional and developmental characterization of cell-type-specific enhancers ^52^. In addition, this approach is applicable to other comparative analyses, such as comparing the different EGI profiles of a cell type between healthy and diseased states, as enhancer activity in certain disease states is altered ^53^.

The ABC model itself is unbiased for distance between enhancers and genes, highlighting its uniqueness as the majority of current methods favour proximity EGIs ^54–56^. Even though, according to previous work, high percentages of EGIs do interact with a neighboring gene ^7,27^, especially for ubiquitously expressed genes where distal enhancers have less effect on their expression, distal EGIs complete the gene expression profile characterization picture. Distal regulations are largely of interest in disease mechanism research, as in disease states such as cancer, long range EGIs could be a contributing factor for development and progression of the disease ^53^.

Last but not least, we have mapped cardiac-phenotype-associated single-nucleotide polymorphisms (SNPs) to the enhancer regions of the EGIs identified in the three cardiac cell types aiming to easy the gene-identification challenge caused by non-coding SNPs. Our methodology is to identify disease-relevant genes through their interactions with enhancers that host cardiac-associated SNPs. To illustrate the validity of our approach, we have been able to identify a QT-interval-associated variant in an endothelical cell-specific enhancer region regulating NOS3 gene expression on chromosome 7, which is due for future experimental validation. This could be the first of many discoveries of identifying a disease-relevant gene modified by a non-coding variant. These results have not only complemented our previous study ^30^, but also raised the significance of SNPs that are in linkage disequilibrium (LD) with sentinel SNPs but have not yet been extensively studied. The previous publication focused on the EGIs of CMs and demonstrated the capability of identifying disease-relevant genes and variants using the ABC model. Our findings have extended this approach to several non-CM cell types and further included genomic variants in linkage equilibrium with the sentinel SNP identify the true causal variant.

A major limitation of our approach is the lack of integration of actual gene expression data into the model. Indeed, in future studies it would be beneficial to validate these results through integration of target gene expression.

As we carried out this study as a proof-of-concept, we acknowledge as a major limitation our reduced sample size and the variation in our CF samples. In future studies and through single-cell analysis it would be important to control for and investigate covariates like age and sex, as well to identify potential subtypes of non-CMs ^45^.

In all, these results, although limited by sample size and covariates, suggest that the described method is a reliable way for linking non-coding GWAS variants with disease-relevant genes through the identification of the interactions between genes and enhancers, the latter of which host these non-coding variants. It has economic benefits and has huge potential to accelerate future disease-relevant gene identification through EGI characterization. This approach can be universally be applied to further disease conditions and cell-types.

## MATERIALS AND METHODS

### Cell lines, sample acquisitions, and library preparation

Human coronary artery smooth muscle cells (HCASMC) and Human coronary artery endothelial cells (HCAEC) were all commercially bought from Thermo Fisher Scientific, Waltham, MA, U.S.A. (product code: C-017-5C) and the American Type Culture Collection (ATCC), Manassas, VA, U.S.A. (Product code: PCS-100-020), respectively. Two of the cardiac fibroblast (CF) samples (*cf_5, cf_6*, Table S2) were from donors who passed away due to non-cardiac diseases and were collected through the National University Hospital (Singapore) surgeon collaborations. The rest of the human cardiac fibroblast samples were purchased from Promocell, Heidelberg, Germany (Product code: C-12375) in the exception of *cf_1*, which was purchased from Cell Applications, San Diego, CA, U.S.A. (product code: 306-05f). A breakdown of the sample cell types and their usage in sequencing was summarized in Fig. 2 A:

ChIP-seq and ATAC-seq were performed as previously described ^57–59^. For each ChIP-seq library, cell pellets were rinsed twice with cold PBS. Cells were crosslinked in 1% formaldehyde. Cells were lysed in lysis buffer (50 mM HEPES-KOH, pH 7.5, 150 mM NaCl, 1 mM EDTA, 1% Triton X, 0.1% Sodium deoxycholate, 0.1% SDS, Takara 1X protease inhibitors) using a glass douncer tight pestle A with 10 to 15 strokes to release nuclei. Centrifugation was performed at 4000 rpm for 10 min at 4řC to collect the nuclear pellet. Nuclei were checked under microscopy for each preparation. Nuclei were lysed in nuclei lysis buffer (50 mM HEPES-KOH, pH 7.5, 150 mM NaCl, 1 mM EDTA, 1% Triton X, 0.1% Sodium deoxycholate, 1% SDS, Takara, 1x protease inhibitor) and sonicated with a Bioruptor sonicator to obtain chromatin fragments between 200 to 500 bp. Sheared chromatin was immunoprecipitated with 5 µg of H3K27ac antibody (Abcam, ab4729) with 50 µl of protein G beads (Invitrogen) overnight at 4řC. Beads were washed and eluted in 200 µl of elution buffer (50 mM Tris-HCl, pH 7.5, and 10 mM EDTA), de-crosslinked at 65řC overnight. Pulldown extracts was phenol/chloroform treated, and DNA was purified by ethanol precipitation. Library preparation was performed using 2 ng of ChIP DNA with the NEB Ultra II library preparation kit, according to the manufacturers protocol. 1012 PCR cycles were performed using indexed primers, and libraries of size between 300 to 500 bp were selected to undergo paired-end sequencing at 2 Œ 150 bp read length on Illumina HiSeq 4000.

For ATAC-seq, cells were lysed in an ATAC-resuspension buffer containing 0.5% NP40, 0.5% Tween-20, and 0.01% Digitonin. The nuclei were then resuspended in a 50 µl transposition mixture containing 25 µl of 2x TD buffer, 16.5 µl of PBS, 5 µl nuclease-free water, 0.5 µl of 1% digitonin, 0.5 µl of 10% Tween-20 and 2.5 µl of transposase (Illumina Tagment DNA enzyme 1, #20034198) and incubated at 37řC for 30 minutes with agitation. DNA was extracted using the NEB Monarchő PCR & DNA cleanup kit (#T1030L) and used for PCR using the Illumina/Nextera primers. Library clean-up was performed using Ampure XP beads and sequenced on a NextSeq (Illumina).

HiChIP was performed using the Arima Hi-C Kit (Arima Genomics, Carlsbad, CA, USA) according to the manufacturer’s protocol. For the initial ChIP steps, the nuclear pellet was sonicated to obtain chromatin fragments between 200 to 500 bp. Sheared chromatin was immunoprecipitated with 3 µg of H3K27ac antibody (Abcam, ab4729) with 30 µl of protein G beads(Invitrogen) overnight at 4žC. Beads were washed and eluted in 200 µl of elution buffer (50 mM sodium bicarbonate, 1% SDS), de-crosslinked at 67žC for 2 hours, and DNA extracted using the Qiagen MinElute kit (Cat no. 28004) according to manufacturer’s instructions. Following to the manufacturer’s protocol, library preparation was performed using the NEB Ultra II library preparation kit (Product no. NEB E7103). Amplification was performed for 10-12 PCR cycles were performed using indexed primers, and libraries of size between 300 to 500 bp were selected. The libraries underwent paired-end sequencing at 2 × 151 bp read length on Illumina HiSeq 4000.

### H3K27ac ChIP-seq and ATAC-seq data processing

Sequencing reads were processed using the according pipelines established by the nf-core community ^60^. Briefly, sequencing reads were first quality-checked and trimmed off the adapter sequences using FastQC ^61^ and Cutadapt ^62^, respectively. Subsequently, reads were aligned and indexed using BWA ^63^, the Homo sapiens genome assembly GRCh38 (hg38) was used as the reference genome. PCR duplicates and multiple-mapping reads were filtered out using SAMtools ^64^. Peaks were called using MACS2^65^ with a threshold of 0.1. Peaks overlapping with blacklist regions ^66^ were filtered out. Reads in peaks were counted using featureCounts ^67^. The quality control step is included in the corresponding nf-core pipelines run by MultiQC ^68^, where metrics such as the number of reads, duplicated reads percentage, percentage of reads mapped, and Fraction of Reads in Peaks (FRiP) are evaluated as shown in supplementary table S2. Since the library preparation process for the chromatin conformation capture HiChIP demands a higher number of cells, to still maintain the instructed number of cells for ChIP-seq, similar samples have been pooled, and HCAECs have been excluded for differential analysis due to a lack of replicates.

Due to the low sample size of this proof-of-concept study, we decided to exclude sex chromosomes from the analysis in order to filter out positives caused by sex as a covariate.

### HiChIP data processing

H3K27ac HiChIP libraries were first processed by the nf-core/hic pipeline. In short, after quality control using FastQC ^61^, HiC-Pro software, which includes alignment software bowtie2^69^, was used to generate contact maps ^70^. Normalized contact maps at 5kb resolution were created using the software Cooler ^71^.

A brief summary of software usage has been listed in the following table:

**Table 1:** Bioinformatic software used for each preprocessing step.

| Software used | ChIP-seq | ATAC-seq | HiChIP | Description |
| --- | --- | --- | --- | --- |
| FastQC<sup>61</sup> | Yes | Yes | Yes | Quality check after sequencing. |
| Cutadapt<sup>62</sup> | Yes | Yes | Yes | Adapter Trimming. |
| BWA<sup>63</sup> | Yes | Yes | No (Bowtie2<sup>69</sup>) | Reads mapping. |
| MACS2<sup>65</sup> | Yes | Yes | No | Peak Calling. |
| Hi-C pro<sup>70</sup> | No | No | Yes | DNA interaction identification and contact matrix creation. |
| MultiQC<sup>68</sup> | Yes | Yes | Yes | After processing quality check. |

### Differential chromatin landscape analysis

R package Diffbind ^34^ was used to identify differential ATAC-seq peaks and differential H3K27ac ChIP-seq peaks between cell types. Heatmap and PCA were produced, using the appropriate functions in the package, to verify sample correlations and similarities. Regions that were in the ENCODE blacklist were removed ^66^. Only significantly differential peaks that have been reported by both DESeq2^72^ and edgeR ^73^ methods, which were dependencies of Diffbind, are reported to limit false positive rates. Differential analysis was only carried out where at least two data sets from distinct biological replicates per cell type where available. Thus, no differential analysis was carried out on HiChIP data or HCAEC ChIP-seq. A false discovery rate (FDR) less than 0.05 and the absolute value of log2 fold change greater than 1 were used as thresholds to identify statistically significant differential peaks. Differential peaks were annotated using the R package ChIPseeker ^74^.

### Enhancer-gene interaction computation

Enhancer-gene interactions (EGIs) were computed using the Activity-by-contact (ABC) model according to the authors’ instructions ^27^, adding a few supplementary processing steps to accord for our specific datasets. Before calling peaks, sex chromosomes were manually removed to exclude possible noise from sex chromosomes. Then peaks were called using MACS2^65^ with summit identification parameters (*–call-summits -p 0*.*1*). Subsequently, one consensus peak set was generated for each cell type to offset variations introduced by sample-specific peaks, using Diffbind. This consensus peak set was used for the next step, enhancer activity (A) estimation. The strength of enhancers was measured by normalizing the number of ATAC reads within and H3K27ac ChIP reads around the putative enhancer regions. The ATAC peaks could also suggest whether a gene from the gene pool is activated or not (genomic coordinates from the ABC Git repository). Then, EGIs were defined using the putative enhancer list, gene list, and contact maps, which were estimated from 3D chromatin architecture (HiChIP). The normalized contact maps from the nf-core/hic pipeline were modified so that the file format is aligned with the ABC model required input format for chromatin conformation capture data, using the R package HiCcompare ^75^. Finally, ABC scores for each EGI are calculated according to the following formula 1, where estimated enhancer activity (A) is multiplied by contact frequency (C) between the two DNA segments from chromatin conformation data before being normalized against all EGIs targeting that gene.

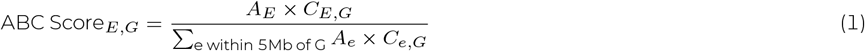

### Common EGI identification and significant threshold

Pairwise comparison of EGIs were carried out between cell-types. Common EGIs were defined as those EGIs that appeared in both cell types compared, allowing a 50% mismatch for the enhancer region (roughly 250 bp). The bedtools software (v2.30.0) ^76^ was used for this processing step applying the following parameters: *-wa -wb -f 0*.*5 -r*, followed by a target gene verification step, where the targetGene were matched. Since bedtools only looked for region overlaps, a target gene verification step was performed after, which culminates in all the common EGIs. To identify those EGIs that indicated a highly cell-type-specific acivity, we used the common EGIs from the pariwise comparisons to compute a delta ABC score using R. A threshold specific to each delta ABC distribution (mean of all the delta ABC *±* 5 *×* the standard deviation (SD)) was selected (See supplementary fig. S2).

In the ABC model, a enhancer is redefined as ‘self-promoters’ if it is less than 750 away from its target gene and is forcefully assigned a high ABC score of 1 (See supplementary fig. S3). We noticed that some enhancers were labeled as ‘self Promoters’ in one cell type, but not in the other cell types (See supplementary fig. S4) because we allowed allowed for a 50% overlap of the enhancers. This led to the EGIs in different cell types were identified as the same, but the ‘self-promoter’ label were different. To exclude them from affecting the delta ABC calculation and differential EGI identification, we adapted the code such that the enhancer regions are consistent for ‘self-promoter’ label.

### Cell-type-specific EGI filtering

EGI filtering was carried out through 2 major steps. First, target genes that existed only in the gene list of one cell type and not the others, regardless of the enhancers, those EGIs were automatically considered as cell-type-specific. For the rest of the EGIs, if the enhancer regions from different cell types, while targeting the same genes, overlapped with each other less than 50% of their enhancer length, we considered those EGIs cell-type-specific. Finally, to define biologically relevant cell-type-specific EGIs, we used a threshold for the ABC score greater than 0.05 as suggested in the original publication for reporting.

### Mapping of published cardiac-disease-associated GWAS signals to enhancers of cell-type-specific EGIs

Publicly available data from various genome-wide association study (GWAS) investigating cardiac traits and/or disease states (hypertrophic and dilated cardiomyopathy, Q-T interval, arrythmia, atrial fibrillation, cardiac arrest, ventricular ectopies) were obtained from the GWAS Catalog ^35^. Phenotypes from GWAS catalogue were categorized into broader categories of hypertrophic and dilated cardiomyopathies, Q-T interval variability, arrhythmias, atrial fibrillations, sudden cardiac arrests, and Duchenne muscular dystrophy. About 1,000 SNPs were associated with one or more cardiac phenotypes. To account for genomic regions in linkage disequilibrium (LD), we further included more than 67,000 SNPs that are in LD with the 1,000 sentinel SNPs (LD *r*^2^ > 0.6; LD between GWAS sentinel SNPs and LD SNPs was calculated using plink software ^77^). Using bedtools, we subsequently mapped those SNPs to the enhancer regions of the previously identified EGIs.

### Version Information

**Table 2:** Version information of software used

| Software | Version |
| --- | --- |
| **nf-core/chipseq** | 2.0.0 |
| **nf-core/atacseq** | 2.0 |
| **nf-core/hic** | 2.1.0 |
| **Nextflow** | 22.10.8 |
| **fastqc**<sup>61</sup> | 0.11.9 |
| **cutadapt**<sup>62</sup> | 3.4 |
| **bwa**<sup>63</sup> | 0.7.17-r1188 |
| **SAMtools**<sup>64</sup> | 1.15.1 and 1.16.1 |
| **macs2**<sup>65</sup> | 2.2.7.1 |
| **featureCounts**<sup>67</sup> | 2.0.1 |
| **MultiQC**<sup>68</sup> | 1.13 and 1.14 |
| **Hi-C pro**<sup>70</sup> | 3.1.0 |
| **bowtie2**<sup>69</sup> | 2.4.4 |
| **cooler**<sup>71</sup> | 0.8.11 |
| **Diffbind**<sup>34</sup> | 3.4.11 |
| **DESeq2**<sup>72</sup> | 1.34.0 |
| **edgeR**<sup>73</sup> | 3.36.0 |
| **R** | 4.1.2 |
| **ChIPseeker**<sup>74</sup> | 1.30.3 |
| **Activity-by-contact (ABC)**<sup>27</sup> | 1.0.0 |
| **HiCcompare**<sup>75</sup> | 1.16.0 |
| **bedtools**<sup>76</sup> | 2.30.0 |
| **Python** | 3.7 |
| **plink**<sup>77</sup> | 1.9 |

## AUTHOR CONTRIBUTIONS

Conceptualization and study design: C.G.A., J.H., and Y.Z., Data collection: C.G.A., Data analysis and figures: Y.Z., Bioinformatic supports: JC.G., R.P., O.T., and CJM.L., Initial draft writing: Y.Z., Draft edition: All co-authors.

## DATA AVAILABILITY

All sequencing data related to this manuscript will be deposited in a public repository.

---

**Table S1:** Sex of ATAC-samples.

| Sample ID | Sex |
| --- | --- |
| cf_1 | F |
| cf_2 | M |
| cf_3 | M |
| cf_4 | F |
| cf_5 | M |
| cf_6 | M |
| ec_1 | M |
| ec_2 | M |
| ec_3 | M |
| smc_1 | F |
| smc_2 | F |

**Table S2:** Sequencing stastistics for ATAC-seq

| Sample ID | Total Library (M) | Mapped (M) | Duplicates (M) | Uniquely Mapped percentage | readL | Percentage reads in peaks | Total peaks |
| --- | --- | --- | --- | --- | --- | --- | --- |
| cf_1 | 77.283406 | 77.005468 | 13.233423 | 82.517125% | 135.378091 | 69.2174% | 117051 |
| cf_2 | 80.248162 | 79.762082 | 11.406938 | 85.1797% | 136.550584 | 65.9348% | 113471 |
| cf_3 | 75.871916 | 75.474181 | 11.164087 | 84.761394% | 141.089255 | 65.3978% | 121600 |
| cf_4 | 81.350898 | 80.948914 | 12.11876 | 84.608966% | 141.598797 | 71.7215% | 106543 |
| cf_5 | 82.147616 | 81.677887 | 12.439838 | 84.284916% | 139.357035 | 68.9144% | 115180 |
| cf_6 | 74.035588 | 73.607319 | 9.751151 | 86.250639% | 138.062538 | 66.3667% | 114630 |
| ec_1 | 79.251584 | 78.984576 | 38.69951 | 50.831875% | 125.791892 | 63.425% | 65328 |
| ec_2 | 76.961656 | 76.712207 | 33.666567 | 55.931281% | 125.938289 | 66.2488% | 68483 |
| ec_3 | 74.92718 | 74.609043 | 8.028536 | 88.860287% | 124.244873 | 46.8415% | 81096 |
| smc_1 | 78.242018 | 77.930593 | 28.264565 | 63.477437% | 127.922463 | 72.2668% | 95729 |
| smc_2 | 74.044994 | 73.737999 | 26.545284 | 63.735186% | 127.951425 | 70.4945% | 91515 |

**Figure S1:**
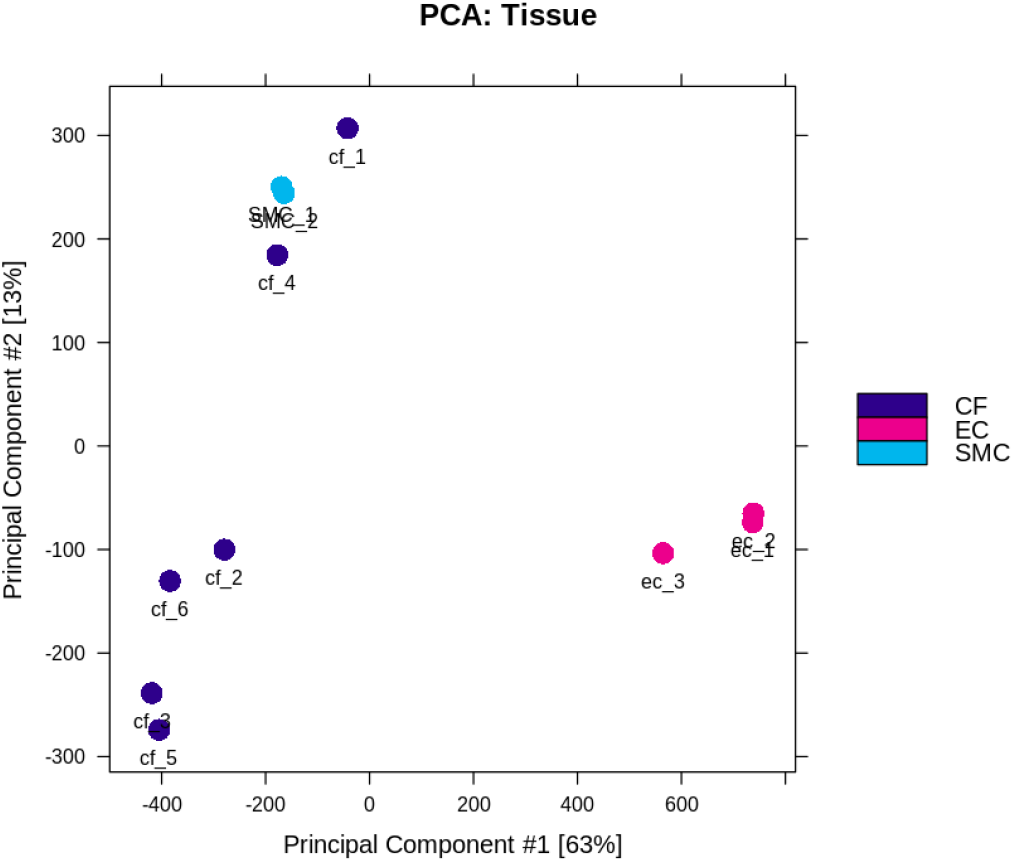
ATAC-seq principal component analysis (PCA). To determine whether chromatin accessibility patterns differed between the three cardiac cell types, principal component analysis (PCA) and correlation heatmaps (Fig 2B) were applied to ATAC-seq peak calls across all samples. The analysis showed that each cell type mostly clustered with samples from the same cell type. There are larger variations between CF samples after we have retraced the samples, the differences have been acknowledged to age and sex differences (Table S1).

**Figure S2:**
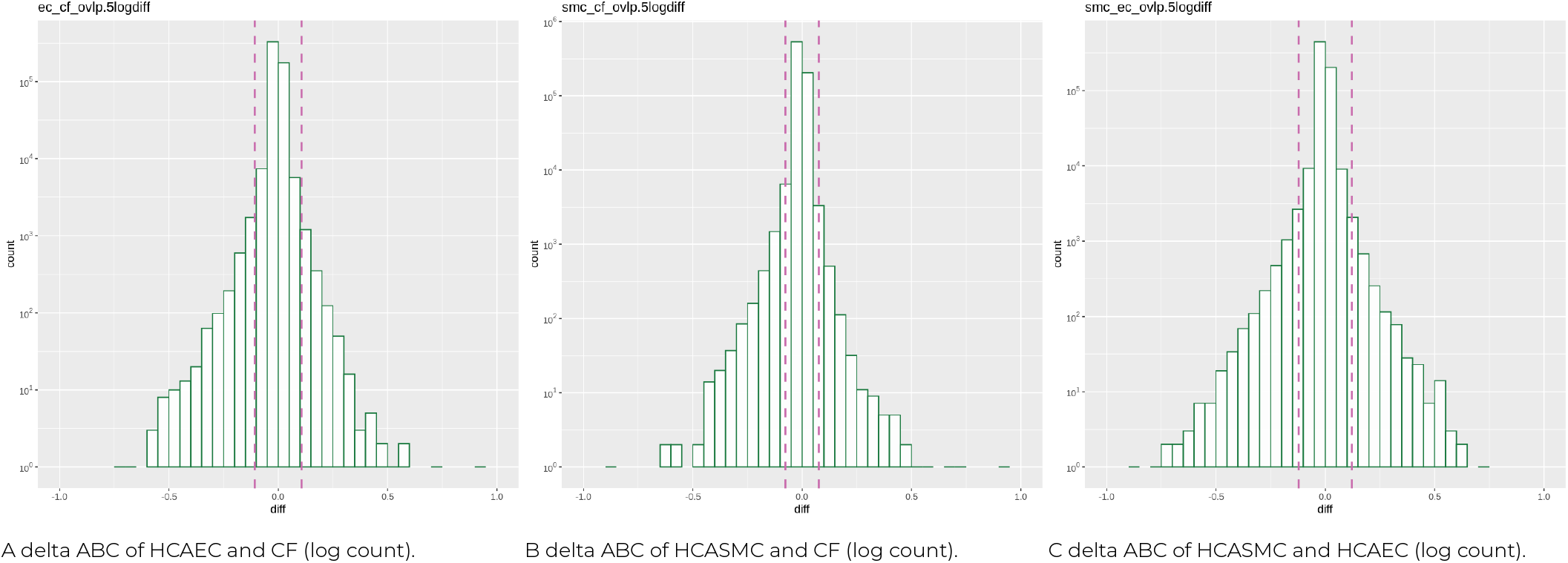
Delta ABC and reasoning for defining significant threshold. Visualization of the delta ABC score distribution with a log scale. We noticed that the majority of delta ABCs are near zero, which was in the center. The pink vertical line represents mean *±* 5 *×* standard deviation (SD). (a) Positive: the EGI has higher score in EC. (b) Positive: the EGI has higher score in SMC. (c) Positive: the EGI has higher score in SMC.

**Table S3:** Number of putative enhancers identified by ABC. After looking for co-localization of H3K27ac histone modification and ATAC-seq (open chromatin) signals, the model identified 133,834, 79,227, and 94,590 putative enhancers for CF, HCASMC, and HCAEC, respectively.

| CF | HCASMC | HCAEC |
| --- | --- | --- |
| 133,834 | 79,227 | 94,590 |

**Figure S3:**
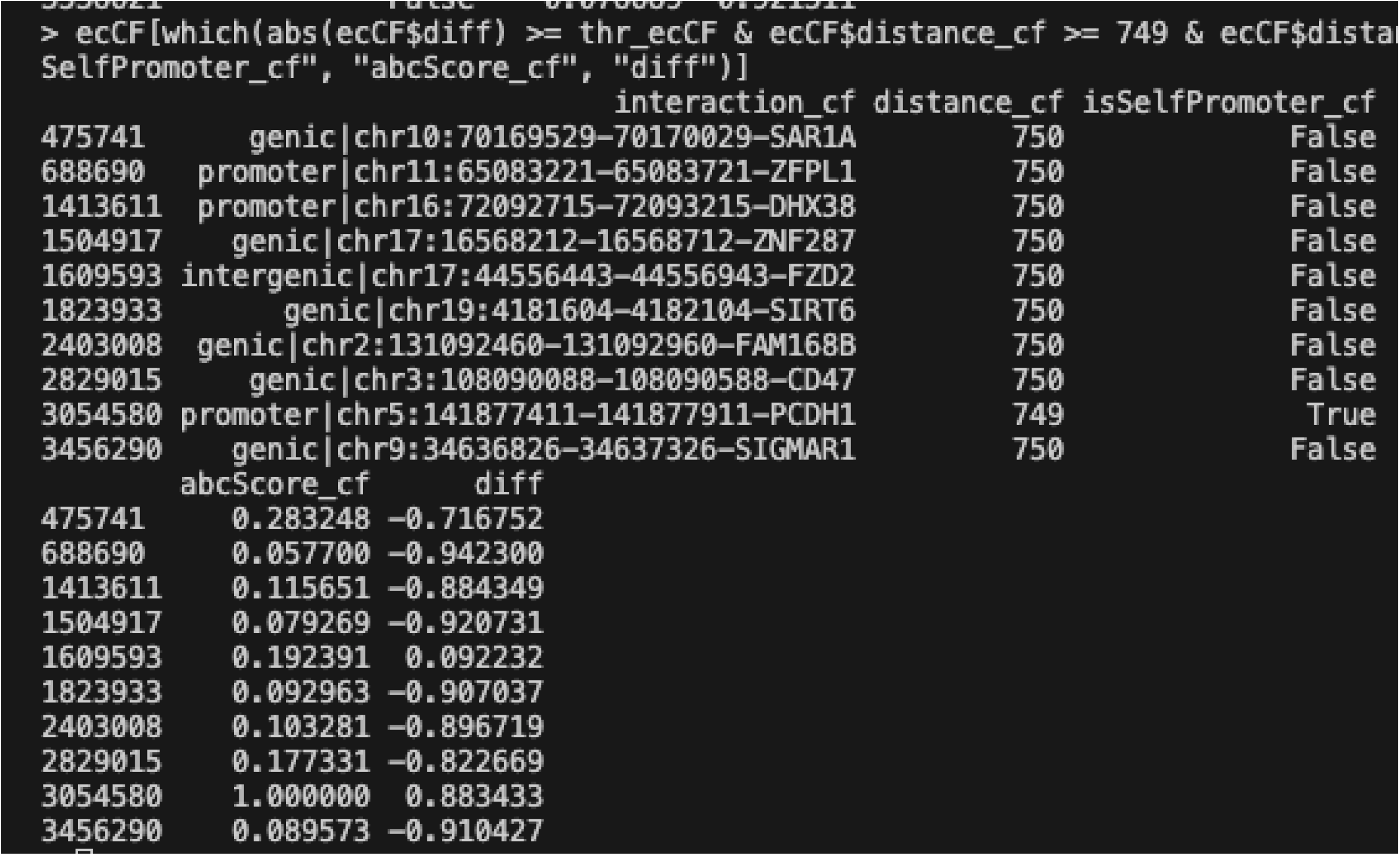
Only putative elements less than 750 bp away from the target gene will be defined as self-promoters. We realized that the model defines promoter based on a hard cut-off of less than 750 bp from the gene. Albeit this will capture the majority of promoters, exceptions could exist as some promoters lie further upstream ^78^. In our analysis, if one element was defined as a promoter in one cell type, we argue that the nearby region in the other cell type should also be an enhancer, as only a 250 bp mismatch was allowed.

**Figure S4:**
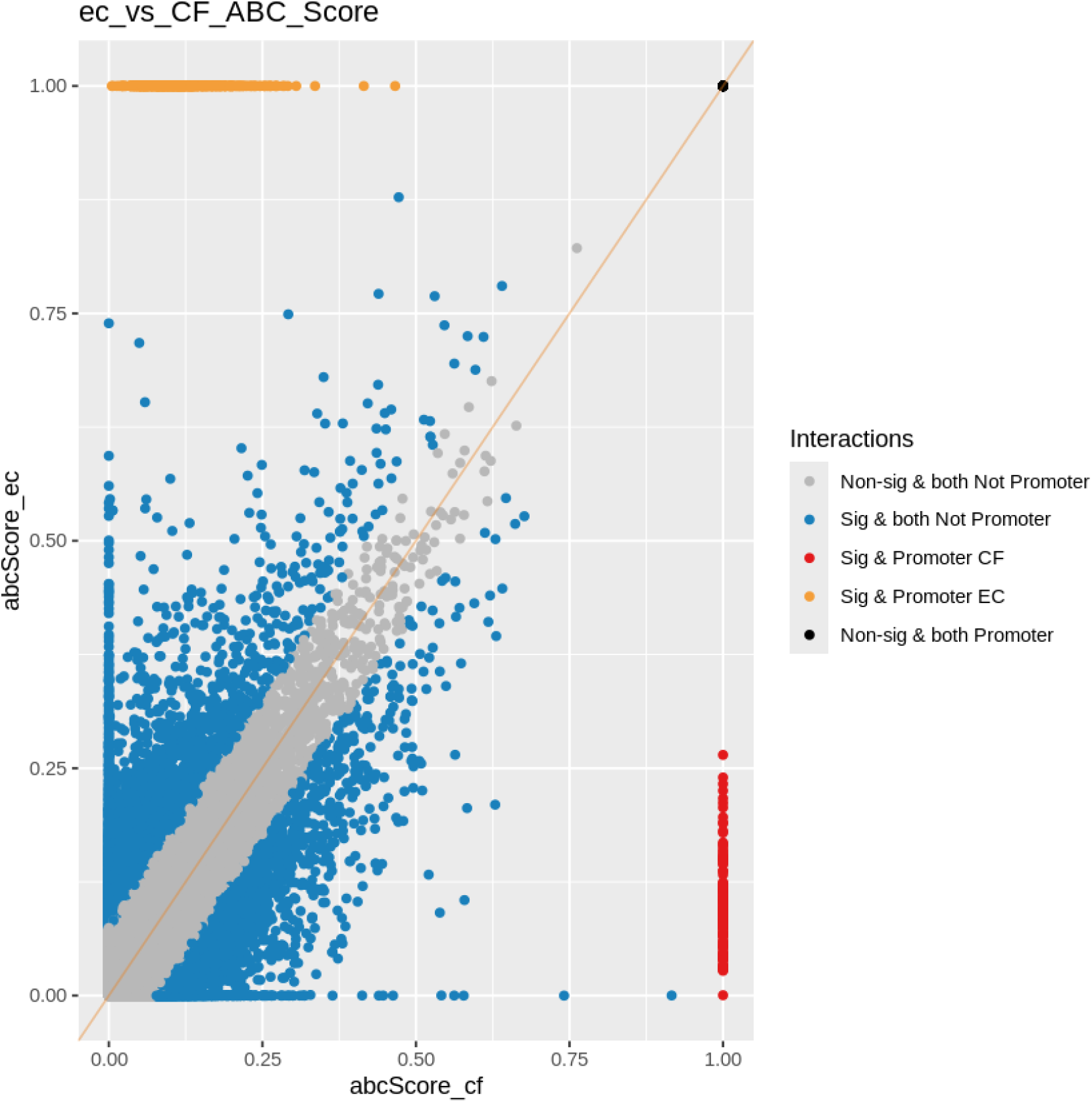
Pair-wise common enhancer-gene interactions (EGIs) and abc score distribution using unadapted model. The black, grey, and blue class EGIs are the same as fig 3. For the yellow and the red groups, those particular enhancers have been defined as self-promoters in one cell type, but not in the other cell type.

**Figure S5:**
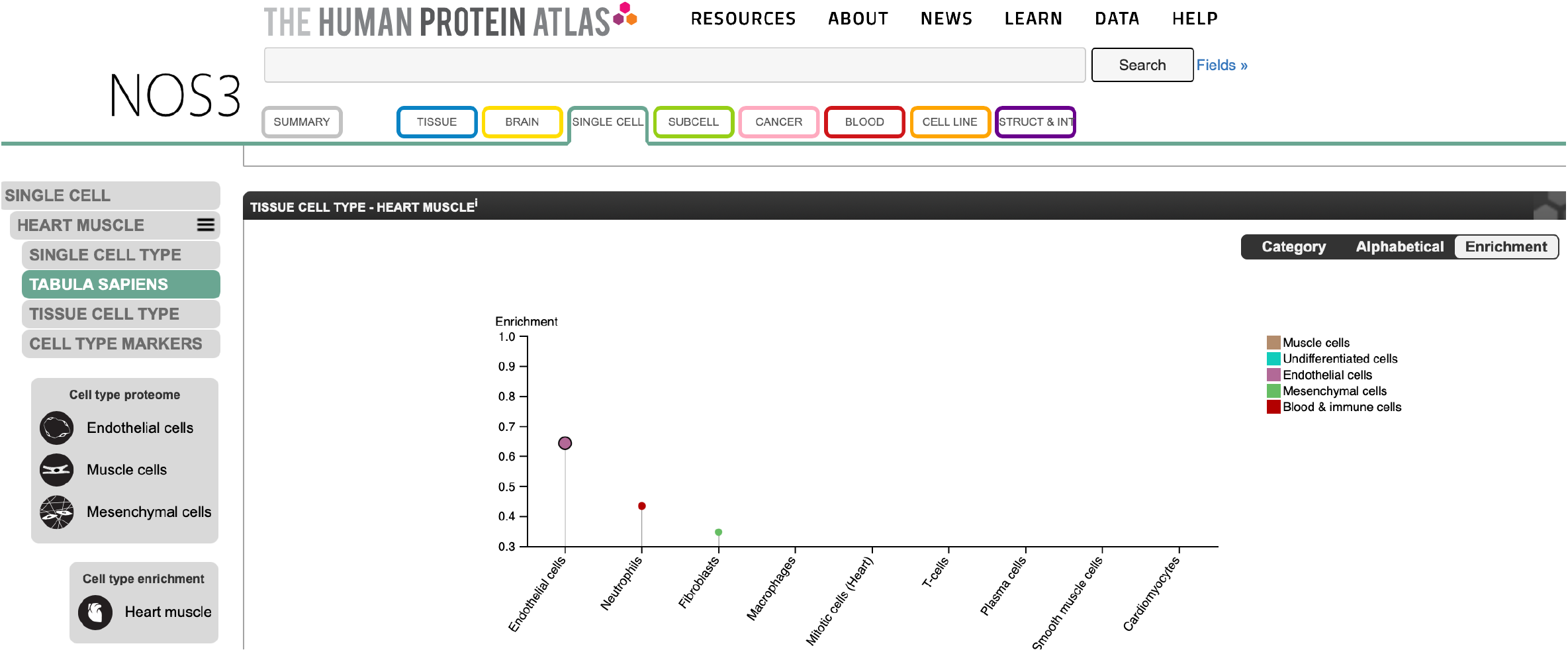
NOS3 expression in various cardiac cells according to The Human Protein Atlas ^36^. The highest enrichment is in endothelial cells.

**Figure S6:**
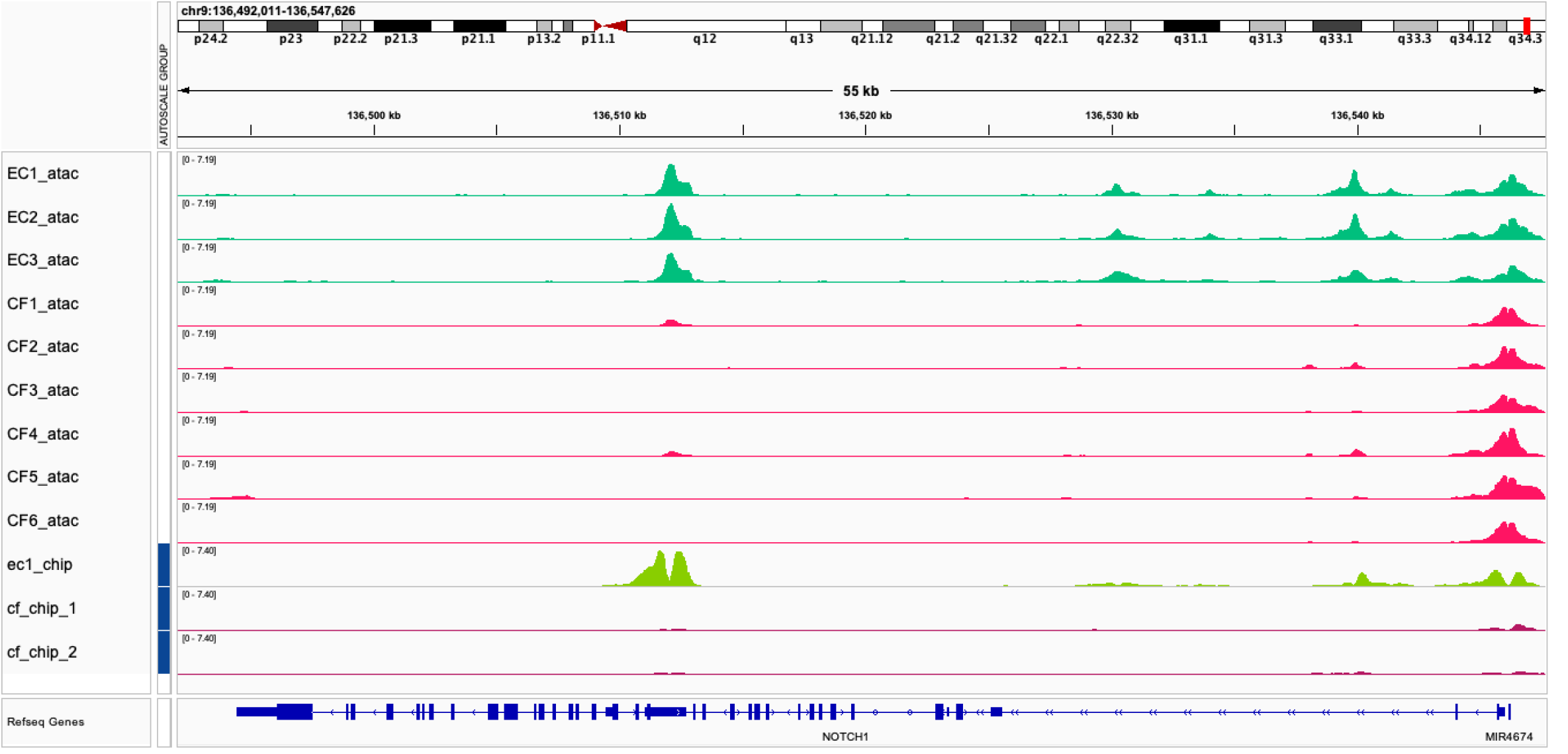
NOTCH1 has an EC-specific ATAC-peak.

**Figure S7:**
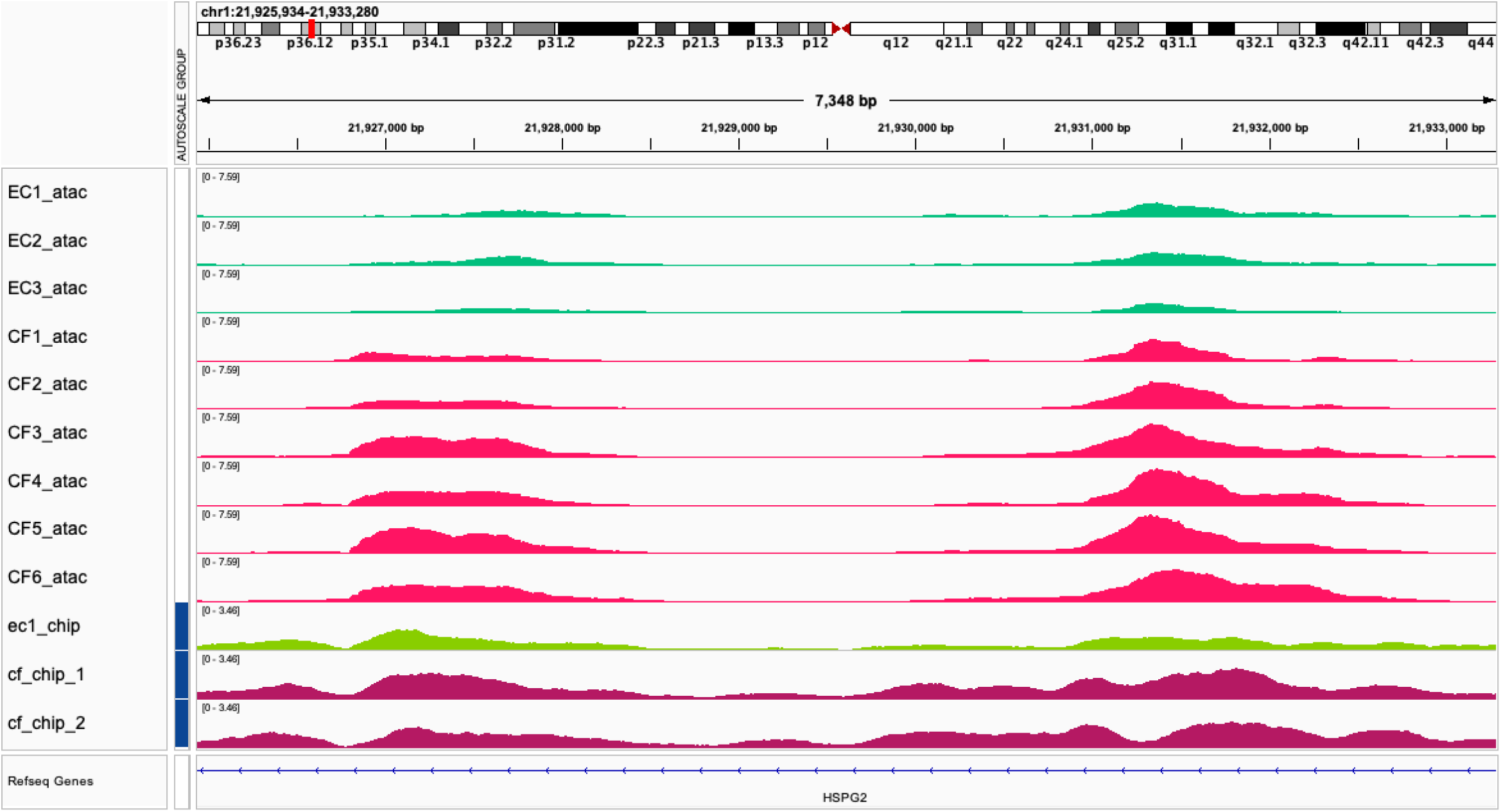
CF-specific ATAC-peaks of HSPG2.

